# Modulation effect of non-invasive transcranial ultrasound stimulation in an ADHD rat model

**DOI:** 10.1101/2022.09.12.507687

**Authors:** Mengran Wang, Teng Wang, Hui Ji, Jiaqing Yan, Xingran Wang, Xiangjian Zhang, Xin Li, Yi Yuan

**Author notes:** Corresponding authors: Xin Li, No.438, Hebei Street, Qinhuangdao 066004, China. Yi Yuan, No.438, Hebei Street, Qinhuangdao 066004, China.

## Abstract

Previous studies have demonstrated that transcranial ultrasound stimulation (TUS) with noninvasive high penetration and high spatial resolution has an effective neuromodulatory effect on neurological diseases. Attention deficit hyperactivity disorder (ADHD) is a persistent neurodevelopmental disorder that severely affects child health. However, the neuromodulatory effects of TUS on ADHD have not been reported to date. To reveal the neuromodulatory effects, TUS was performed in ADHD model rats for 2 consecutive weeks, and the behavioral improvement of ADHD, neural activity of ADHD from neurons and neural oscillation levels, and the plasma membrane dopamine transporter (DAT) and brain-derived neurotrophic factor (BDNF) in the brains of ADHD rats were evaluated. The results showed that TUS can improve cognitive behavior in ADHD rats, and that TUS altered neuronal firing patterns and modulated the relative power and sample entropy of local field potentials in an ADHD rat model. In addition, TUS can also enhance BDNF expression in the brain tissue of the ADHD rat model. In conclusion, TUS has an effective neuromodulatory effect on ADHD and thus has the potential to clinically improve cognitive dysfunction in ADHD.

## Introduction

Childhood attention deficit hyperactivity disorder (ADHD), commonly known as hyperactivity disorder, is a chronic brain disorder characterized by inattention, hyperactivity, and impulsivity(Association, 2013). ADHD is prevalent in 2%–7% of children and adolescents globally, with an average prevalence rate of 5%(Association, 2013; Sayal, Prasad, Daley, Ford, & Coghill, 2018). Children diagnosed with ADHD typically have functional impairment in multiple domains, and more than half of them have ADHD comorbidities such as anxiety and depression (Danielson et al., 2018; Liang, Zou, Cen, & Li, 2009). In addition, ADHD is a typical long-term behavioral disorder, and its damage will continue into adulthood (Hinshaw, 2018) and even lead to conduct disorder, adult ADHD, and juvenile delinquency. This seriously affects the mental health of school-age children, causes harm to individuals, and places a heavy burden on families and society. Therefore, it is important to choose an appropriate diagnosis and treatment plan according to the specific symptoms of children with ADHD.

Drug therapy is the most widely used clinical treatment for ADHD. Commonly used drugs include central nervous system stimulants (methylphenidate hydrochloride) and central norepinephrine-regulating drugs (atomoxetine), with central nervous system stimulants being recommended as the first-line treatment for ADHD. Drug treatment can achieve good results, have a faster effect on children with severe symptoms, and relieve the symptoms of attention deficit and hyperactivity in children(D. R. Coghill et al., 2014). However, drug treatment has side effects such as loss of appetite, sleep disturbance, temporary growth retardation in height, cramps, and substance abuse(Ahn, Ahn, & Cheong, 2016). Non-drug physical interventions are thus included in the treatment of ADHD. Behavioral intervention and neurofeedback training have been shown to have significant effects on reducing ADHD symptoms and behavioral problems(D. Coghill, 2010; Knopf, 2021). Transcranial direct current stimulation (tDCS) and transcranial magnetic stimulation (TMS) have become popular and promising physical therapy techniques in recent years owing to their advantages of convenience and high safety(Bikson et al., 2016). Studies have shown that tDCS can modulate behavioral, redox, and inflammatory parameters in animal models of ADHD and as well as significantly improve attention, inhibitory control, and working memory(Cosmo et al., 2020; Dubreuil-Vall et al., 2021; Leffa et al., 2018), and its efficacy is closely related to stimulation parameters. A previous study demonstrated that TMS can modulate neuronal excitability and alter brain electrical activity, thereby improving the clinical symptoms of ADHD patients(Alyagon et al., 2020).

In addition to the aforementioned physical therapy techniques, non-invasive transcranial ultrasound stimulation (TUS), a new neuromodulation technology, has developed rapidly in recent years. Ultrasound waves are used to penetrate the entire brain and skull to achieve noninvasive neuromodulation. Compared with tDCS and TMS, TUS has finer spatial specificity and a higher penetration depth(Bystritsky et al., 2011; Landhuis, 2017; Meng, Hynynen, & Lipsman, 2021; Yu, Niu, & He, 2020). Previous studies have evaluated the modulatory effect of TUS on healthy mice with respect to behavior, neural activity, cerebral hemodynamics, and protein expression. These studies found that TUS could induce the motor response of the samples, enhance or inhibit the synchrony of neuronal firing and related neural network activity, modulate cerebral hemodynamics, and increase the expression of neurotrophic factors (NTFs)(H. Kim, A. Chiu, S. D. Lee, K. Fischer, & S.-S. Yoo, 2014; Liu, Lai, Chen, & Yang, 2017; Yusuf Tufail et al., 2010; X. Wang, Yan, Wang, Li, & Yuan, 2019; X. Wang, Zhang, Zhang, & Yuan, 2021; Yoo et al., 2011; Yu, Niu, Krook-Magnuson, & He, 2021; Yu, Sohrabpour, & He, 2016; Yuan, Wang, Liu, & Shoham, 2020). These studies have shown that TUS has a significant modulatory effect on neural activity and its corresponding neurovascular coupling. With respect to potential clinical applications, ultrasound stimulation is helpful in the treatment of neurological diseases, including epilepsy.

Ultrasound stimulation can suppress the number of epileptic signal bursts, reduce seizure duration, increase the interval between seizures, and reduce the power spectrum intensity of low frequencies of neural activity(Li et al., 2019; Z. Lin et al., 2020; Min et al., 2011; Zou et al., 2020) In Parkinson’s disease, ultrasound stimulation improved behaviors such as pole climbing and forced swimming and significantly reduced correlation in mice models of Parkinson’s disease. Ultrasound stimulation can also protect neural firing activity and protect dopamine neurons from damage by downregulating Bax expression and upregulating Bcl-2 expression, blocking mitochondrial release of cytochrome c (Z. Wang, Yan, Wang, Yuan, & Li, 2020; Yuan, Zhao, et al., 2020; Zhou et al., 2019). In Alzheimer’s disease, ultrasound stimulation effectively improved spatial learning and memory ability in mice models of Alzheimer’s disease (AD). It also significantly improved neuropsychological scores for up to 3 months in AD patients and increased the levels of brain-derived neurotrophic factor (BDNF) and glial cell line-derived neurotrophic factor (GDNF) in AD rat brain (Beisteiner et al., 2020; W.-T. Lin, Chen, Lu, Liu, & Yang, 2015; Popescu, Pernet, & Beisteiner, 2021; Zhang et al., 2022). These findings demonstrate that TUS has significant therapeutic and rehabilitative effects, such as improving behavior, modulating neural firing activity, and increasing/decreasing disease-related protein expression, in various neurological diseases. Therefore, it is a potential alternative for the clinical treatment of neurological diseases. However, it is unclear whether TUS has an effective therapeutic effect on ADHD, a neurological disorder.

Children with ADHD have symptoms of inattention, difficulty in motor control, impulse suppression, and behavior regulation (Roberts, Milich, & Barkley, 2015), which seriously affect their daily lives and social communication(Miller et al., 2018). This attention-deficit, hyperactive-impulsive behavior in children with ADHD is considered an external manifestation of specific brain regulatory dysfunction(Weyandt & Gudmundsdottir, 2014). In addition to behavioral abnormalities, patients with ADHD also have abnormal neural firing activity, which is mainly reflected in the changes in brain activity on electroencephalography (EEG). Previous studies have shown that children with ADHD have abnormal EEG spectrum in some conventional neuropsychological modes (Hobbs, Clarke, Barry, McCarthy, & Selikowitz, 2007), mainly manifested as increased slow wave activity of EEG and increased absolute and relative power of theta and delta EEG (Fox, Tharp, & Fox, 2005; Hughes & John, 1999; Vincent J Monastra et al., 2006). Fast wave activity decreases, and the absolute and relative power of alpha wave and beta wave decreases (Fonseca et al., 2008; King, Barkley, Delville, & Ferris, 2000; Steven M Snyder & James R Hall, 2006). At the molecular level, neurotrophins are considered important targets in ADHD clinical manifestations and pathogenesis (Galvez-Contreras, Tania, Gonzalez-Castaneda, & Oscar, 2017; Steven M Snyder et al., 2008). In particular, BDNF has been shown to be associated with the pathogenesis of ADHD and may thus serve as a biological target for its treatment(Galvez-Contreras et al., 2017; Tsai, 2017). In addition, dopamine neurotransmitters, especially the plasma membrane dopamine transporter (DAT), also play an important role in ADHD pathogenesis (D. Leo, Sukhanov, Zoratto, Illiano, & Gainetdinov, 2018). However, it remains unclear to date whether TUS modulates ADHD-related physiological indicators, such as behavior, neural firing activity, and protein expression.

Thus, this study aimed to evaluate the modulation effect of non-invasive TUS in ADHD. Towards this goal, we performed TUS in ADHD rat models for 2 consecutive weeks. Behavioral improvements were evaluated, and then neuronal action potentials and local field potentials were recorded on days 1, 4, 7, 14, and 21 after stimulation initiation. The effect of TUS on ADHD rat models neural activity from neurons and neural oscillation levels was investigated. Finally, the changes in DAT and BDNF in the brain tissue of ADHD rat models under ultrasound stimulation were studied.

## Results

### Effects of low-intensity transcranial ultrasound stimulation on cognitive dysfunction

Various behavioral experiments were performed to study the regulatory effect of ultrasonic stimulation on cognitive function in an ADHD rat model. In the open field experiment, compared with the Wistar Kyoto rat (WKY) group and the Sprague-Dawley (SD) group, the spontaneous hypertension (SHR) group showed obvious hyperactivity symptoms and lower SHR hyperactivity symptoms under TUS. Fig. 1a shows the activity trajectories of the rats in each group. To accurately analyze the regulatory effect of ultrasound stimulation in the ADHD rat model, we quantitatively analyzed the movement distance, movement time, and central zone residence time of all rats (Fig. 1c-e). The SHR group was compared with the SD and WKY groups, and the SHR group had longer movement distance (SD group: 1947.0 ±152.9, WKY group:1099.0±149.2, SHR group:6325.0±464.5). Compared with the SHR group, the SHR+TUS group had approximately 35% less movement distance. The total movement time of the SHR group was also significantly longer than that of the other groups (SD group: 144.1±20.67, WKY group: 77.96±13.82, SHR group: 485.6±14.18). The total movement time was also significantly lower by approximately 27% in the SHR+TUS group than that in the SHR group. Further, the area of interest (ROI) is the central area in the open field, the SHR group stayed longer in the ROI (SD group: 2.146±1.157, WKY group: 2.369 ±1.542, SHR group: 80.69±10.96). Similarly, the center dwell time was lower by 46% in the SHR+TUS group than in the SHR group. (Mean ± SEM; n = 6 for each group; *p < 0.05, **p < 0.01, Kruskal–Wallis test)

**Figure 1.**
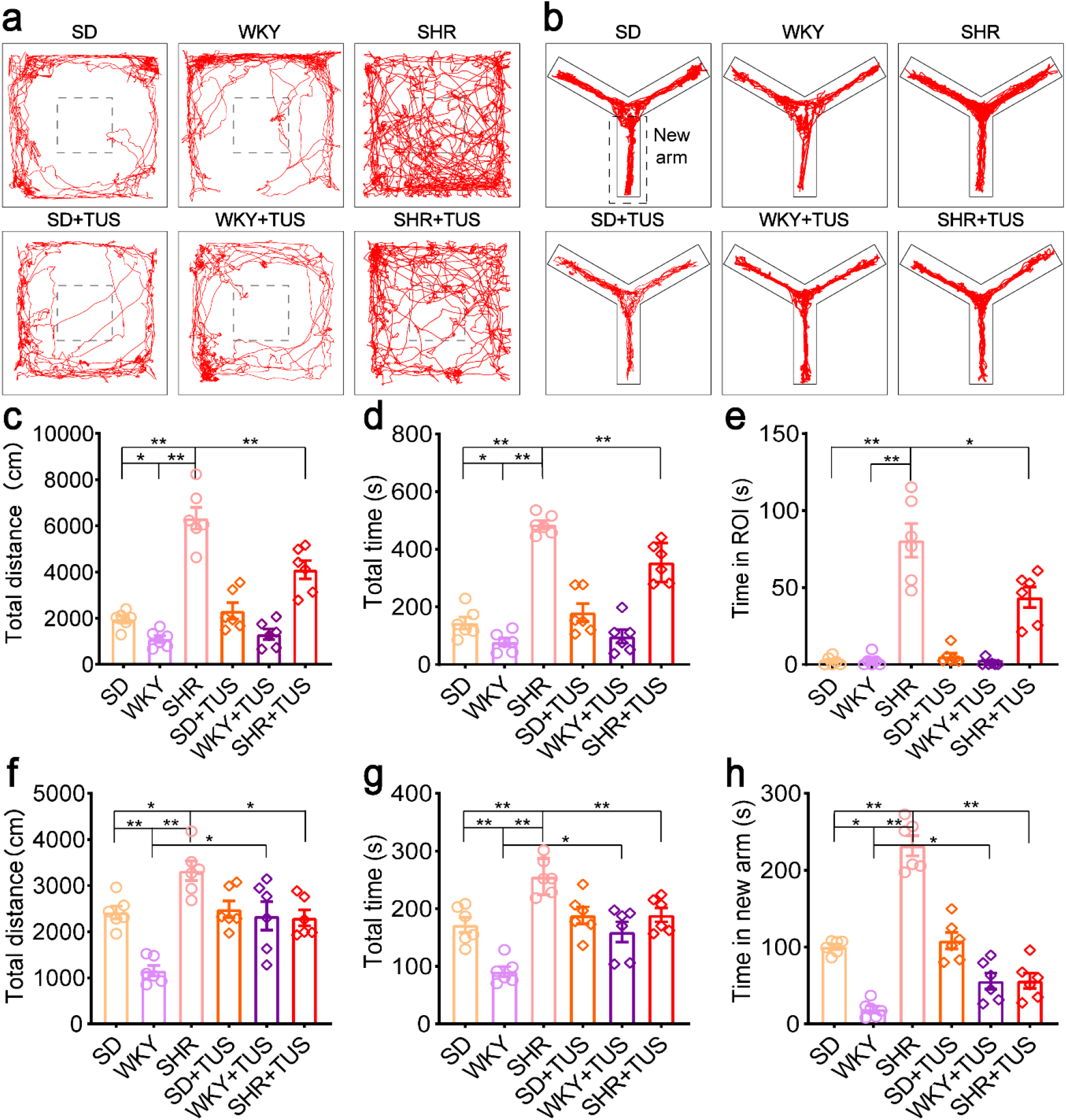
Open field test and Modified Y-maze test. (a)Movement paths taken on the open field test in different groups. (b) Movement paths taken in the modified version of the Y-maze test in different groups. (c)Total distance travelled on the open field test in different groups. (d)Total time travelled on the open field test by rats in different groups. (e)Time in ROI travelled on the open field test by rats in different groups. (Mean±SEM, n=6 for each group in the open field test, *p <0.05, **p<0.01, Kruskal–Wallis test).(f) Modified Y-maze results, total distance travelled in different groups. (g) Modified Y-maze results, total time travelled by rats in different groups. (h) Modified Y-maze results, time in new arm travelled by rats in different groups. (Mean±SEM, n=6 for each group in the modified Y-maze test, *p <0.05, **p<0.01, Kruskal–Wallis test).

The movement trajectories of each group in the modified Y-maze are shown in Fig. 1b. We also quantified the movement distance, movement time, and dwell time in the novel arm for all rats (Fig. 1f–h). Compared to the SHR group, the SHR+TUS group showed lower total distance traveled by approximately 31% and lower exercise time by approximately 26%. The time spent was also significantly reduced in the new arm by approximately 76%. Concurrently, we unexpectedly found that compared with the WKY group, the WKY + TUS group had significantly higher total movement time, total movement distance, and dwell time of the new arm. (Mean ± SEM, n = 6 for each group, *p < 0.05, **p < 0.01, Kruskal–Wallis test).

In the delayed non-matched placement version of the T-maze (Fig. 2a), the accuracy of the SHR group was significantly lower than that of the SD and WKY groups (Fig. 2b). The daily change in the correct rate of the different groups are shown in Fig. 2c-e. The correct rate of the SD, SD+TUS, WKY, and WKY+TUS groups fluctuated, but there was no significant change after 10 days of stimulation. However, changes were more obvious in the SHR+TUS group than in the SHR group. During the 10-day stimulation process, the correct rate of the SHR+TUS group gradually increased from 57.1±5.2% on the first day to 83.3±4.3% on the tenth day (Mean ± SEM, n = 6 for each group, *p < 0.05, **p < 0.01, Kruskal–Wallis test).

**Figure 2.**
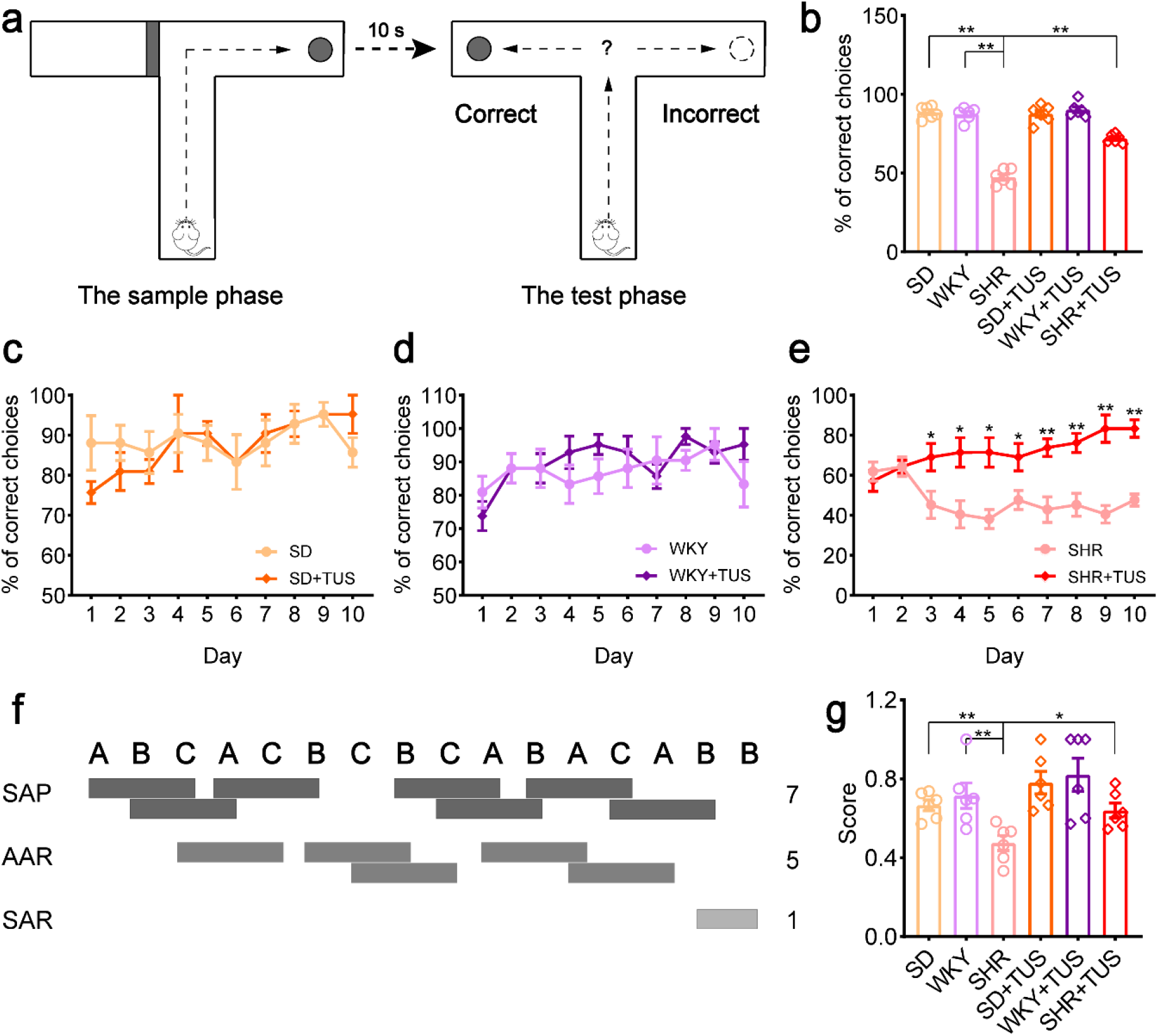
DNMTP T-maze and Y-maze spontaneous alternation task. (a)DNMTP T-maze. (b) A DNMTP of the T-maze results, total correct choices for 10 days. (c) The change of correct choices for 10 days on the T-maze test in SD and SD+TUS groups. (d) The change of correct choices for 10 days on the T-maze test in the WKY and the WKY+TUS groups. (e) The change of % of correct choices for 10 days on the T-maze test in the SHR and the SHR+TUS groups. (Mean±SEM, n=6 for each group in DNMTP T-maze, *p <0.05, **p<0.01, Kruskal–Wallis test).(f) Performance of Y-maze autonomous alternation task. (g) alternation performance of SAP on Y-maze spontaneous alternation task. (Mean±SEM, n=6 for each group in Y-maze spontaneous alternation task, *p <0.05, **p < 0.01, Kruskal–Wallis test).

Performance of Y-maze autonomous alternation task was shown in Fig.2f. The SHR+TUS group exhibited significantly higher spontaneous alternation than did the SHR group (Fig. 2g) (Mean ± SEM, n = 6 for each group, *p < 0.05, **p < 0.01, Kruskal–Wallis test), but there was no significant change between other control groups and its ultrasound stimulation groups (Mean ± SEM, n = 6 for each group, *p > 0.05, Kruskal–Wallis test). These behavioral test results indicate that ultrasonic stimulation can affect behavior and alleviate cognitive dysfunction in ADHD rats.

### Effects of low-intensity transcranial ultrasound stimulation on neural activity

We used ultrasound to stimulate the prefrontal cortex of rats in each group and recorded neural activity using multichannel electrodes to evaluate neuronal spikes (Fig. 3a). The spikes were classified using Boss software, the interference was removed, and the firing of each neuron was finally obtained (Fig. 3b-c). The firing rate of neuronal action potentials was then analyzed. To quantitatively assess changes in the firing rate, the firing rate of neuronal spikes was calculated in each group. As shown in Fig. 3d–f, the firing rates of the neuronal spikes in the SD and WKY groups did not change significantly after ultrasound stimulation. In contrast, the firing rate of neurons in the SHR group were significantly increased on days 7, 14, and 21 after ultrasound stimulation (Mean ± SEM, n = 6 for each group, *p < 0.05, **p < 0.01, Kruskal–Wallis test). The daily neuronal firing rate in each group was determined, and the SHR group showed a downward trend within 21 days and an upward trend after ultrasound stimulation (Figure supplement 1). The SHR group had lower neuronal spiking rates on days 1, 4, 7, and 21 than did the SD and WKY group (Figure supplement 2). These results suggest that ultrasound stimulation can modulate neural activity in SHR by increasing the firing rate of spikes in SHR neurons.

**Figure 3.**
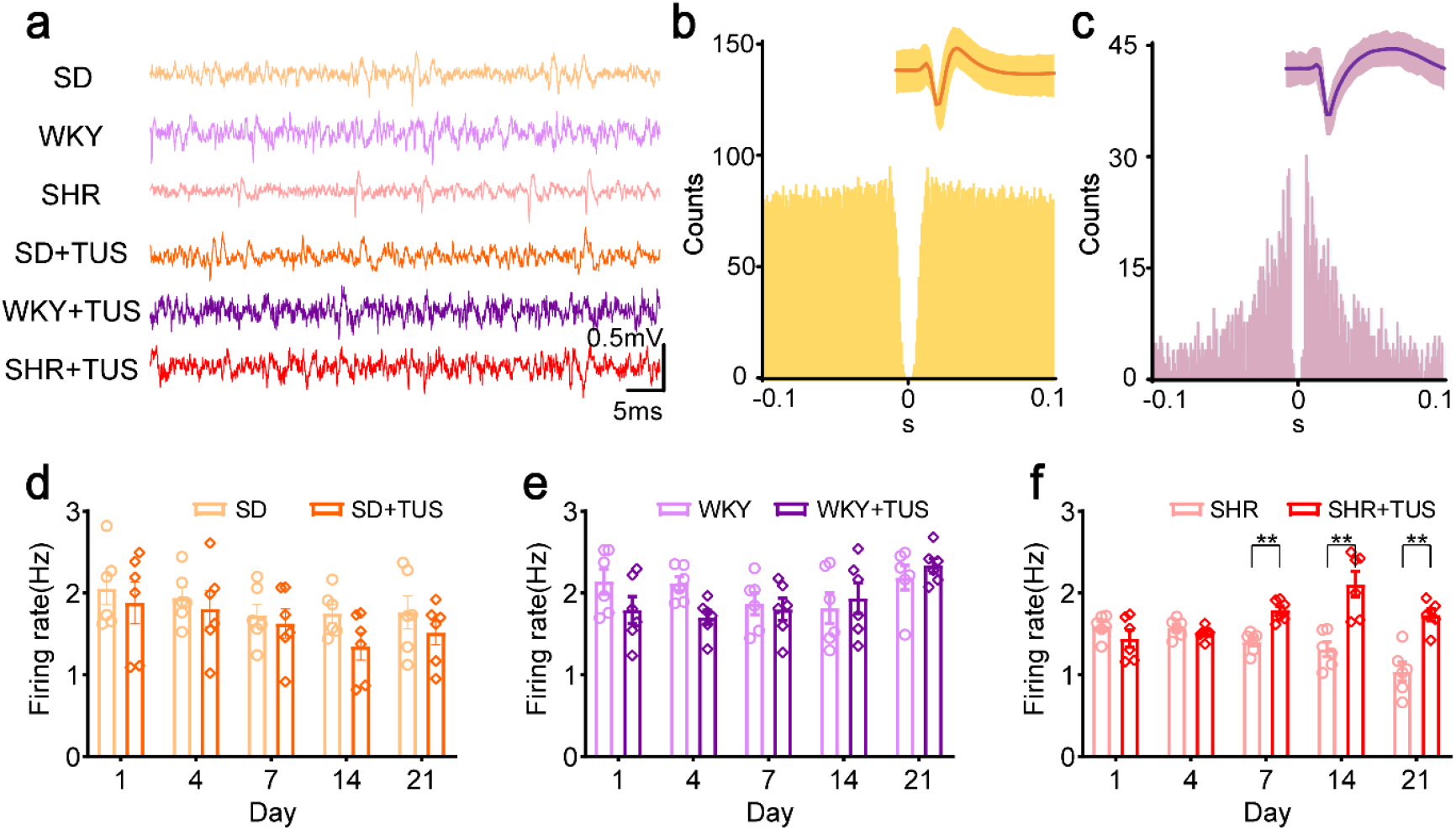
High-pass signal and average of spike firing rate. (a) High-pass signal (> 300Hz) of different groups. (b) The autocorrelation plot of an interneuron. (c) The autocorrelation plot of a pyramidal neuron. (d)-(f)Average of spike firing rate of all neurons in different groups, (d) SD and SD+TUS groups, (e) WKY and WKY+TUS groups, (f) SHR and SHR+TUS groups. (Mean±SEM, n=6 for each group, **p<0.01, Kruskal– Wallis test).

Based on the study of neuronal spiking rates, the changes in neural oscillations caused by ultrasound stimulation was further evaluated. The relative power and sample entropy of theta, beta, and gamma frequency bands of the local field potential (Fig. 4a) were then analyzed. The relative power of the theta, beta, and gamma frequency bands in the SD+TUS group did not differ significantly compared with that in the SD group (Fig. 4b–d). Meanwhile, compared to the WKY group, the WKY + TUS group showed significantly higher relative power of the theta band 21 days after the initiation of ultrasound stimulation (Fig. 4e), and the relative power of the beta band was significantly lower on days 14 and 21 of the initiation of ultrasound stimulation (Fig. 4f). Further, the relative power of the gamma band was significantly lower on day 21 after the initiation of ultrasound stimulation (Fig. 4g) (Mean ± SEM, n = 6 for each group, **p < 0.01, Kruskal–Wallis test).

**Figure 4.**
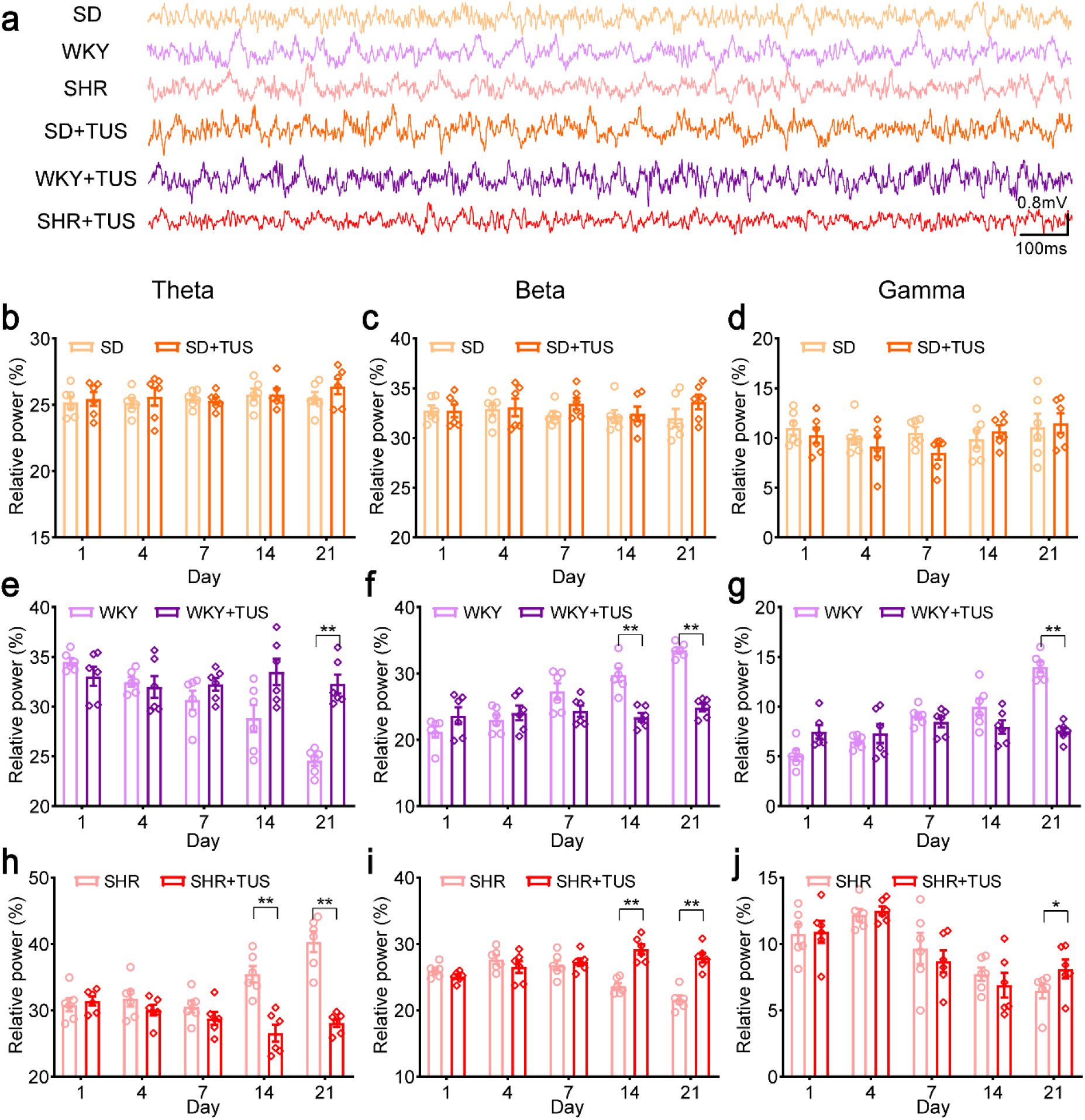
LFPs and the relative power at the different frequency bands. (a) LFPs in the different groups. (b)-(d) The relative power of LFPs at the theta (4–8 Hz), beta (13–30Hz) and gamma (30–45Hz) frequency bands in SD and SD+TUS groups, (b) theta (4–8 Hz), (c) beta (13–30Hz), (d) gamma (30–45Hz). (e)-(g) The relative power of LFPs at the theta (4–8 Hz), beta (13–30Hz) and gamma (30–45Hz) frequency bands in WKY and WKY+TUS groups, (e) theta (4–8 Hz), (f) beta (13–30Hz), (g) gamma (30–45Hz). (h)-(j) The relative power of LFPs at the theta (4–8 Hz), beta (13–30Hz) and gamma (30–45Hz) frequency bands in SHR and SHR+TUS groups, (h) theta (4–8 Hz), (i) beta (13–30Hz), (j) gamma (30–45Hz). (Mean±SEM, n=6 for each group, **p<0.01, Kruskal–Wallis test).

The relative power of theta bands in the SHR+TUS group was significantly lower by 25% and 30% on days 14 and 21 after the initiation of ultrasound stimulation, respectively, compared with that in the SHR group (Fig. 4h). The relative power in the beta band was higher by 24% and 29% on days 14 and 21 after ultrasound stimulation (Fig. 4i), respectively, and the relative power in the gamma band significantly higher by 25% on day 21 after the initiation of ultrasound stimulation. (Fig. 4j) (Mean ± SEM, n= 6 for each group, *p < 0.05, **p < 0.01, Kruskal–Wallis test). In addition, we investigated the relative power of each group of animals as a function of time within the stimulation cycle, where the relative power of the SD, SD+TUS and WKY+TUS groups did not change significantly. The relative power of different frequency bands in the SHR+TUS group showed a significant trend with stimulation time (Figure supplement 3). And we found that the relative power of the SHR group was likely to be significantly different from the SD and WKY groups during the same stimulation time (Figure supplement 4).

Regarding the nonlinear kinetic analysis, the results showed that sample entropy was not significantly changed before and after ultrasound stimulation in the SD (Fig. 5a–c) and WKY groups (Fig. 5d–f). However, compared to the SHR group, the SHR+TUS group showed significantly higher sample entropy in the theta band (Fig. 5g) and significantly lower sample entropy in the beta band (Fig. 5h) on days 4, 7, 14, and 21 after ultrasound stimulation. The sample entropy in the gamma band was significantly lower on day 21 after ultrasound stimulation (Fig. 5i) (Mean ± SEM, n = 6 for each group, *p < 0.05, **p < 0.01, Kruskal–Wallis test). Similarly, we studied the change of the sample entropy of each group of animals during the stimulation period. The sample entropy of SD, WKY, and SD+TUS groups did not change significantly, and the WKY+TUS group showed a trend of change in theta frequency band. The sample entropy of theta-band and beta-band in the SHR+TUS group tended to decrease with increasing stimulation time (Figure supplement 5). And we found that the relative power of SHR group may be significantly different from that of SD group and WKY group in theta and beta frequency bands, but not in gamma frequency band during the same stimulation time (Figure supplement 6). These results indicate that ultrasound stimulation can effectively modulate neural oscillations in rat models of ADHD.

**Figure 5.**
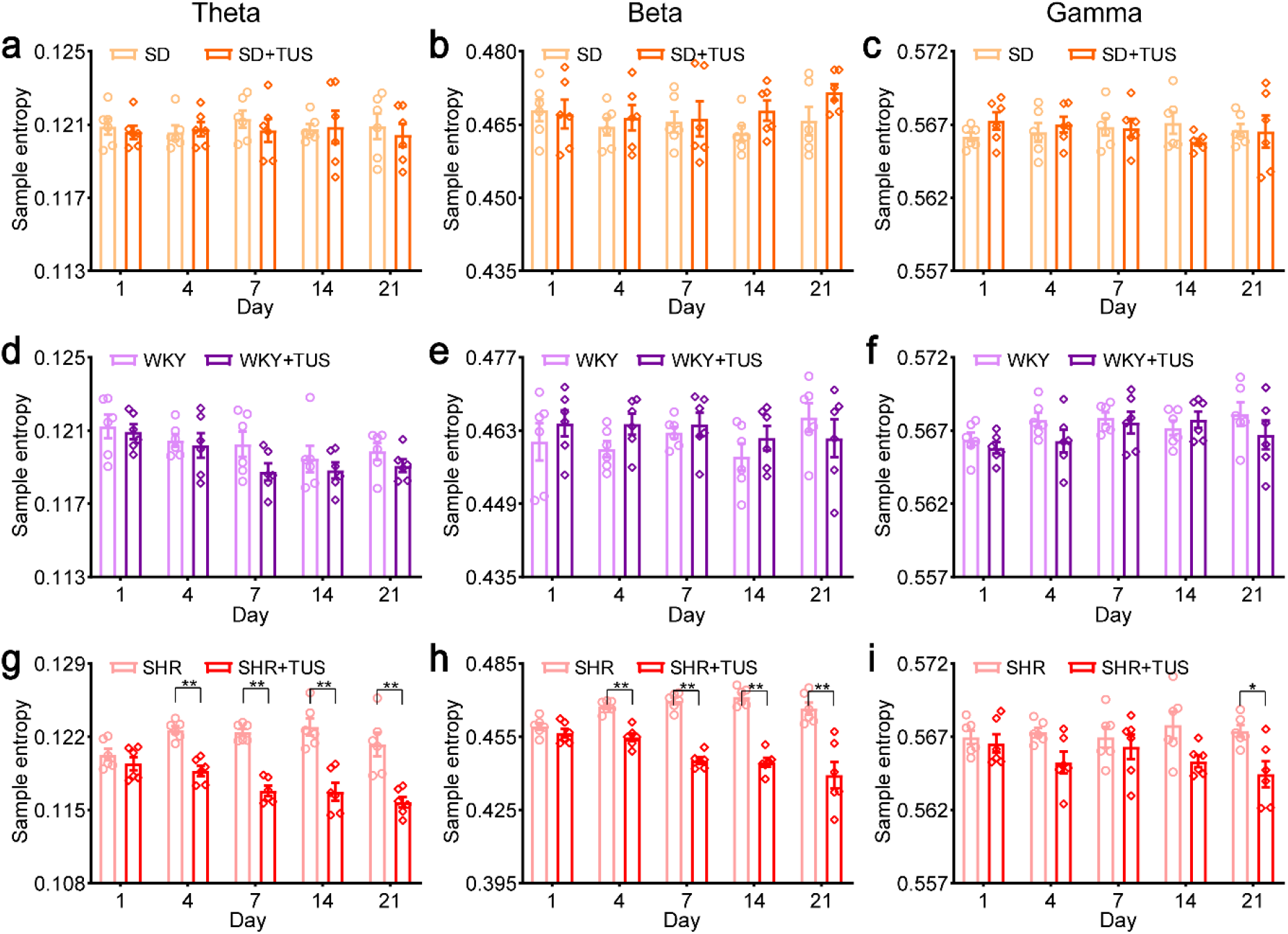
The sample entropy of LFPs at the different frequency band. (a)-(c) The sample entropy of LFPs at the theta (4–8 Hz), beta (13–30Hz) and gamma (30–45Hz) frequency bands in SD and SD+TUS groups, theta (4–8 Hz), (b) beta (13–30Hz), (c) gamma (30–45Hz). (d)-(f) The sample entropy of LFPs at the theta (4–8 Hz), beta (13–30Hz) and gamma (30–45Hz) frequency bands in WKY and WKY+TUS groups, (d) theta (4–8 Hz), (e) beta (13–30Hz), (f) gamma (30–45Hz). (g)-(i) The relative power analysis of LFPs at the theta (4–8 Hz), beta (13–30Hz) and gamma (30–45Hz) frequency bands in SHR and SHR+TUS groups, (g) theta (4–8 Hz), (h) beta (13–30Hz), (i) gamma (30–45Hz). (Mean±SEM, n=6 for each group, *p <0.05, **p<0.01, Kruskal–Wallis test).

### Effects of low-intensity transcranial ultrasound stimulation on the protein expression of DAT and BDNF

Western blot analysis was performed to confirm the changes in protein expression of DAT and BDNF after ultrasound stimulation (Fig. 6a). Compared to the SHR group, the SHR+TUS group showed no significant change in the expression of the DAT after ultrasound stimulation (Fig. 6b). However, BDNF expression was higher by approximately 30% in the SHR+TUS group than in the SHR group (Fig. 6c) (Mean ± SEM, n= 3 for each group, *p < 0.05, Kruskal–Wallis test). These results suggest that ultrasound stimulation of the prefrontal cortex modulates the protein expression of BDNF in a rat model of ADHD.

**Figure 6.**
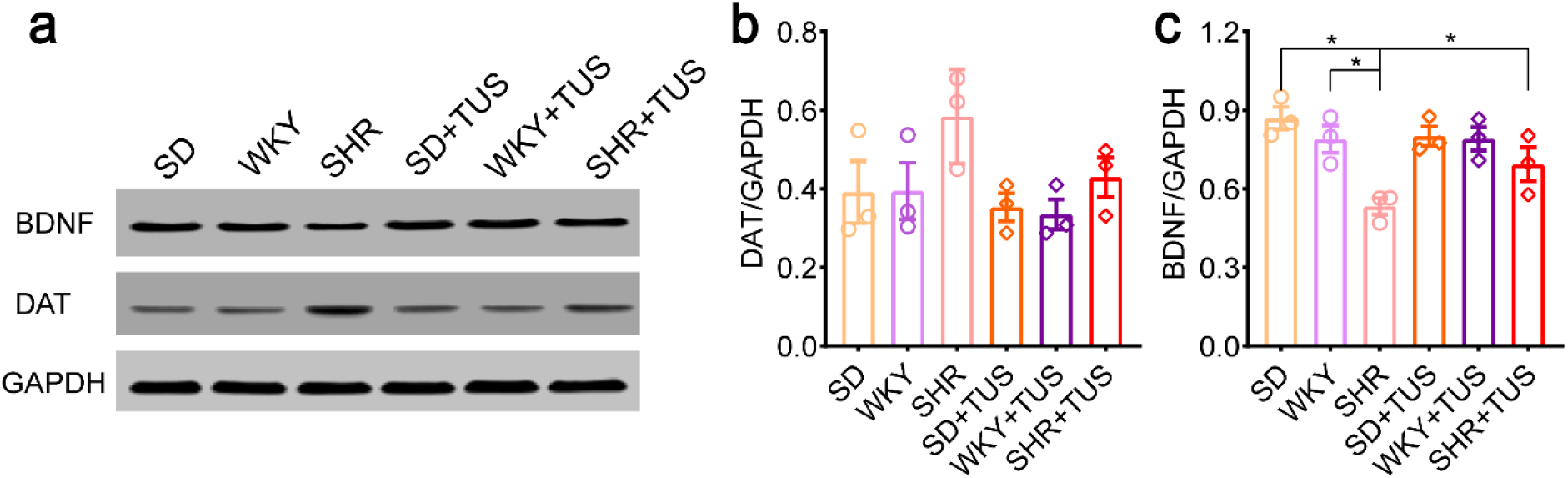
Representative western blots for DAT and BDNF (a) Western blots for BDNF and DAT. (b)-(c) DAT and BDNF levels measured by western blots at 2 days after the last ultrasound stimulation in all groups, (b) DAT levels, (c) BDNF levels (Mean±SEM, n=3 for each group, *p <0.05, Kruskal–Wallis test).

## Discussion

There have been no studies on the neuromodulatory effects of TUS on ADHD to date. The main findings of this study are as follows: TUS (1) can improve cognitive behavior and (2) alters neuronal firing patterns and modulates the relative power and sample entropy of LFPs in an ADHD rat model. TUS (3) can also enhance BDNF expression in the brain tissue of the ADHD rat model. Collectively, these findings suggest that ultrasound stimulation can effectively improve cognitive function in an ADHD rat model. Therefore, TUS is a potential therapeutic tool for treating cognitive dysfunction in ADHD.

In our experiments, TUS stimulation of the prefrontal cortex resulted in similar behavioral effects; reduced hyperactivity; and attenuated cognitive impairments in spatial learning, working memory, behavioral flexibility, and exploratory behavior. EEG studies in ADHD have shown that children with attention-deficit ADHD have increased slow-wave activity and insufficient fast-wave activity (Clarke, Barry, Mccarthy, Selikowitz, & Brown, 2002; Vincent J. Monastra et al., 2006). Meanwhile, the synchronization of cortical neurons is improved, desynchronization is decreased, excitability of neurons is weakened, EEG cycle is prolonged, and frequency is reduced, leading to a slower EEG rhythm and symptoms. Therefore, neuronal firing is an important focus. The results showed that TUS significantly reduced the amplitude of multiple stimulus-evoked potentials(W. Legon et al., 2014)and altered the EEG intrinsic activity and evoked potentials in a frequency-dependent manner(Jma, Wl, Aob, Tfs, & Wjt, 2014). These results suggest that TUS can modulate the neural activity in the brain. In our experiments, ultrasound increased the firing rate of neurons in the prefrontal cortex of SHR rats.

Relative power change is an abnormal EEG characteristic in patients with ADHD. Studies have shown that the slow wave activity of EEG in the frontal cortex of children with ADHD increases, and the absolute and relative power of theta and delta wave EEG also increases. Meanwhile, the fast wave activity decreases, and the absolute and relative power of alpha and beta wave decreases(Corrêa et al., 2008; S. M. Snyder & J. R. Hall, 2006; S. M. Snyder et al., 2008). Studies have concluded that theta rhythm is related to animal behavior; the theta phase is closely related to spatial memory, especially sequential memory; and frontal theta enhancement is prominent in sequential working memory tasks(Hsieh, Ekstrom, & Ranganath, 2011; J & N, 2011). Slow and fast gamma rays may selectively affect memory encoding or memory retrieval at different times(Mably & Lee, 2018). Our study showed that ultrasound stimulation increases the relative power in the beta and gamma bands and decreases the relative power in the theta band in an awake active ADHD rat model. These findings suggest that TUS can increase fast-wave activity in a rat model of ADHD and may help improve cognitive dysfunction in spatial and working memory.

Non-linear kinetic features have been studied for the evaluation of treatment effects (Cerquera, Arns, Buitrago, Gutierrez, & Freund, 2012) and diagnostic analysis (A Fernández et al., 2009) in ADHD. Novel non-linear methods based on entropy are widely used to measure the complexity of EEG signals (A. Fernández, Gómez, Hornero, & López-Ibor, 2013). In this study, the sample entropy in the theta band was significantly increased and that in the beta band was decreased after day 4 of ultrasound stimulation in the ADHD rat model. Many researchers believe that important pathophysiological changes in ADHD are caused by a decrease in central excitability caused by insufficient dopamine secretion (Gornick et al., 2007; Ribasés, Ramos-Quiroga, Hervás, Bosch, & Bayés, 2009). Methylphenidate is the first-line drug used for the clinical treatment of ADHD. It directly or indirectly affects the secretion or reabsorption of dopamine in the central nervous system, thereby increasing the concentration of dopamine in the synaptic cleft and improving the symptoms (Storebø, Ramstad, Krogh, Nilausen, & Gluud, 2015). NTFs, particularly BDNF, are critical for neurodevelopment and plasticity in the adult brain and have been implicated in the pathogenesis of ADHD (Ayhan Bilgiç, Aysun Toker, Ümit IşIk, & İbrahim KIlInç, 2017; Tsai, 2017). The current study found that after 14 days of continuous ultrasound stimulation, the level of BDNF in the prefrontal cortex was significantly increased. This suggests that ultrasound stimulation can improve cognitive dysfunction by modulating BDNF in rat models of ADHD.

The current study found that ultrasound stimulation can improve cognitive impairment; however, the underlying mechanism remains unclear. Using cell electrophysiology and optical labeling techniques, Tyler et al.(Tyler, Tufail, Finsterwald, Tauchmann, & Majestic, 2008) demonstrated that TUS can alter the firing activity of neurons by changing the permeability of the cell membrane sodium, potassium, and calcium ion channels. Importantly, ultrasound can stimulate subcortical circuits, causing neuronal activity and synchronized oscillations(Y. Tufail et al., 2010). It has been established that NTF loss leads to neurodegeneration, whereas exposure to endogenous or exogenous NTFs promotes neuronal survival (LF, 2006). There is also evidence that neurotrophins modulate ion channel activity and expression (Adamson, Reid, & Davis, 2002; Needham, Nayagam, Minter, & OLeary, 2012; Purcell et al., 2013). Previous studies have demonstrated that potassium channels, involved in synaptic excitability and neuroplasticity, may be involved in the pathogenesis of ADHD (Lesch et al., 2008; Mick et al., 2010). Therefore, we speculated that ultrasound stimulation can improve cognitive dysfunction by modulating neuronal activity in ADHD rat models.

Current noninvasive physical techniques for cognitive regulation in ADHD include tDCS and TMS. Previous studies that used tDCS in ADHD rat models found that it improved working memory and behavioral flexibility and alleviated cognitive dysfunction, especially when applied to the prefrontal cortex (Da, Ahn, Pak, Hong, & Choi, 2020). In the current study, ultrasound stimulation of the prefrontal cortex of ADHD rats reduced both working memory impairment and hyperactivity symptoms. This finding is similar to the regulatory role of tDCS. Studies that have applied TMS to the right prefrontal cortex of patients with ADHD have reported significantly lower recording EEG signals, TMS-evoked potentials, and event-related potentials in the stop-signal task in ADHD patients than in matched healthy controls. Event-related potential components were also significantly reduced (Kaskie & Ferrarelli, 2017). In the current study, ultrasound stimulation of the prefrontal cortex of rats with ADHD altered neuronal firing rates and effectively modulated the relative power and sample entropy of EEG-specific frequency bands. Notably, TUS has higher resolution and penetration depth than tDCS and TMS.

A previous study using high-density EEG recordings found that, through the use of EEG coherence, alterations in functional connectivity in children with ADHD could be indicated, with a strong relationship to the prefrontal cortex (Murias, Swanson, & Srinivasan, 2007). Functional magnetic resonance imaging studies have shown that activation of the frontal lobe, cingulate gyrus, and other brain regions is decreased in ADHD children (Bush, Frazier, Rauch, Seidman, & Biederman, 1999). Further, the development and function of the prefrontal cortex of ADHD patients is significantly different from that of normal children, but this difference will be with the development of normal children. It gradually decreases with age(Shaw et al., 2012; Yasumura et al., 2019). Thus, prefrontal structural defects play an important role in the pathogenesis of ADHD and have the potential to serve as early markers of ADHD. Ultrasound, as an emerging neuromodulation modality, has high resolution and stimulation depth. Some studies have applied TUS to the human prefrontal cortex to modulate emotion and the connectivity of the prefrontal cortex emotion-related resting-state network(Sanguinetti, Hameroff, Smith, Sato, & Allen, 2020) and to elicit motor cortex excitability(Gibson et al., 2018). TUS can activate the brain to generate sensory and motor neural responses in the absence of external stimuli(Lee, Chung, Jung, Song, & Yoo, 2016; Lee et al., 2015; A. Leo, Priya, Jerel, & Wynn, 2018). TUS can activate the brain to generate sensory and motor neural responses in the absence of external stimulation. In addition, TUS has inhibitory effects on the primary somatosensory cortex, the unilateral sensory nucleus of the thalamus and the primary motor cortex (Fomenko, Chen, Nankoo, Saravanamuttu, & Chen, 2020; Wynn Legon, Leo, Bansal, & Mueller, 2018; W. Legon et al., 2014). Therefore, it is speculated that ultrasound can regulate abnormal brain regions in ADHD and has thus high potential for the clinical treatment of ADHD.

Previous studies have reported that TUS modulates neural activity through nonspecific auditory responses(Hongsun Guo et al., 2018; Sato, Shapiro, & Tsao, 2018). To evaluate the impact of non-specific auditory responses in TUS, we analyzed the relative power of EEG by stimulating three deaf rats in each group with specific ultrasound according to previous literature(Mohammadjavadi, Ye, Xia, Brown, & Pauly, 2019). As shown in Figure supplement 7, the results were similar between deaf and normal rats. Importantly, these results suggest that the modulating effect of TUS on ADHD was not due to auditory effects. Regarding the thermal effects of ultrasound stimulation, previous studies have shown that the thermal effects on neural activity may be negligible in low-intensity ultrasound stimulation, and most studies have attempted to definitively negate the role of thermal effects in ultrasound-evoked and ultrasound-mediated inhibition(Kim et al., 2012; Naor, Krupa, & Shoham, 2016; SS et al., 2011; Y. Tufail et al., 2010; Wright, Rothwell, & Saffari, 2015). Given that low-intensity ultrasound only produces small temperature changes, these changes are likely to be lower than the heat transfer capability of passive perfusion(Dinno et al., 1989). Further, the low-intensity pulse protocol ensures that the tissue temperature rises less than 0.1°C during sonication(H. Kim, A. Chiu, S. D. Lee, K. Fischer, & S. S. Yoo, 2014; Magee & Davies, 1993). Therefore, the influence of the thermal effect of the ultrasonic stimulation on the experiment is negligible.

Our study has two limitations. First, ultrasound stimulation can improve behavioral impairment in ADHD rat models and alter its neuronal firing and LFP-specific frequency band characteristics. However, it is not known whether ultrasound can improve the dynamic changes in EEG signatures during working memory tasks in rats. In future studies, we will record the EEG of rats during behavioral experiments to investigate the regulatory effects of ultrasound on the behavioral EEG characteristics of ADHD rat models. Second, previous studies have shown that ultrasound parameters can have differential modulatory effects on neural activity(Ye, Brown, & Pauly, 2016; Yuan Yi, Jiaqing, Zhitao, & Xiaoli, 2016). Only one parameter was used in the experiments, and it was unclear whether the chosen parameter was optimal. To determine the optimal parameters for the modulation of ADHD, we will conduct a multiparameter study to determine the ultrasound parameters that have the best therapeutic effect.

## Materials and methods

### Animals and groups

A total of 30 Wistar Kyoto rats (WKY), 30 spontaneously hypertensive rats (SHR), and 30 sprague-dawley rats (SD) (all male, postnatal day 25±1) weighing 100–140 g (Beijing Vital River Laboratory Animal Technology Co., Ltd. China) were used in the experiments. The animals were randomly divided into three groups: behavioral test, LFP acquisition, and biochemistry (Table 1). The animals were housed in standard cages with room temperature control (22 ± 1 °C), ventilation, and a 12-h light-dark cycle. The animals had free access to food and water. All procedures were performed in accordance with the guidelines of the Animal Ethics and Management Committee of Yanshan University and all applicable international, national, and/or institutional guidelines for the care and use of animals.

**Table 1.** Animal groups and numbers.

|  | **Behavioral Test** | **LFP acquisition** | **Biochemical detection** |
| --- | --- | --- | --- |
| **TUS** | Wistar Kyoto rats (n=6) | Wistar Kyoto rats (n=6) | Wistar Kyoto rats (n=3) |
|  | Spontaneously Hypertensive rats (n=6) | Spontaneously Hypertensive rats (n=6) | Spontaneously Hypertensive rats (n=3) |
|  | Sprague-Dawley rats (n=6) | Sprague-Dawley rats (n=6) | Sprague-Dawley rats (n=3) |
| **Control (CTRL)** | Wistar Kyoto rats (n=6) | Wistar Kyoto rats (n=6) | Wistar Kyoto rats (n=3) |
|  | Spontaneously Hypertensive rats (n=6) | Spontaneously Hypertensive rats (n=6) | Spontaneously Hypertensive rats (n=3) |
|  | Sprague-Dawley rats (n=6) | Sprague-Dawley rats (n=6) | Sprague-Dawley rats (n=3) |

### Experimental procedures and surgery

Behavioral tests of cognitive function (n = 6 rats/group) were performed according to the experimental schedule proposed in Figure supplement 8f. All rats were anesthetized with 2% isoflurane before electrode implantation surgery. The skull was shaved, and the skin was cleaned with saline (0.9% sodium chloride). The scalp was then cut along the midline of the skull, and the subcutaneous tissue and periosteum were removed. A 16-channel electrode (4 × 4, Beijing Creation Technologies Co., Ltd. China) was implanted into the prefrontal cortex (1.25 mm left and right, 4.35 mm anterior from the bregma) at a depth of approximately 2.5 mm. The specific location is shown in the Figure supplement 8e. A 5-mm plastic tube with a beveled surface was placed near the electrode implantation site on the opposite side to reserve a space for subsequent ultrasound stimulation. Two more holes were drilled to place the stainless-steel screws for fixation. One screw was connected to the reference electrode of the 16-channel electrode at a position in the cerebellum. Finally, the 16-channel and ground electrodes were completely wrapped and fixed on the skull using dental cement. After surgery, rats were kept in a single cage and allowed to recover for 7 days. Thereafter, ultrasonic stimulation and signal recording were performed.

### Low-intensity transcranial ultrasound stimulation protocol

This study also used the TUS system used in our previous study (Y. Yi et al., 2018). The schematic diagram of ultrasound stimulation is shown in Figure supplement 8a. The pulse signal was generated using a signal generator ((AFG3022C, Tektronix, USA) and amplified using an amplifier (E&I 240 L, ENI Inc., USA) to drive an ultrasound transducer (V301-SU, Olympus, USA) to generate ultrasound waves. A 3D printed conical collimator with an ultrasound coupling gel was attached to the rat head to allow ultrasound to penetrate the rat brain. The hole at the bottom of the conical collimator was 4 mm in diameter. As shown in Figure supplement 8b, the ultrasound signals were a fundamental frequency, pulse repetition frequency, duty cycle, and stimulation duration of 500 kHz, 1 Hz, 5%, and 50 ms, respectively. Two-dimensional ultrasound distribution (Figure supplement 8c) in the x-y plane from the position that is 5 mm below the bottom of the collimator were measured by a calibrated needle-type hydrophone (HNR500, Onda, USA). The reconstruction profile was placed along the dotted white lines in the x–y plane (Figure supplement 8d). The diameter of the focal area measured at the full width at half maximum (FWHM) was ∼5.9 mm. The maximum ultrasound pressure was 0.53MPa and the corresponding spatial peak and pulse-average intensity (I_sppa_) values were 9.36 W/cm^2^, respectively. The hair on the rat head was shaved before TUS and the collimator was sown or tilted in the small hole reserved beforehand to ensure alignment with the stimulation site. The stimulation site was in the prefrontal cortex (1.25 mm left and right, −4.5 mm from bregma). The timing diagram of stimulation and detection was shown in Figure supplement 8f. Ultrasound stimulation was performed continuously for 2 weeks, and the total stimulation time was 15 min per time.

### LFP and spike acquisition

The electrodes on the rat head were connected via flexible wires to a preamplifier, which delivered broadband (0.3–7.5 kHz) neural signals (16 bits at 30 kHz) from the electrodes to a multi-channel data acquisition system software (Apollo, Bio-Signal Technologies, McKinney, TX, USA). Peaks were extracted using a high-pass filter (300 Hz). The LFP signal was sampled at 1 kHz, and the peak and raw data were sampled at 30 kHz. Data were collected on days 1, 4, 7, 14, and 21 after stimulation (Figure supplement 8f).

### Open-field test

The open-field test was used to assess spontaneous motor behavior. The rats were placed in a black plexiglass square box (70 × 70 × 40 cm^3^) that was divided into central (30 × 30 cm^2^) and peripheral areas. Rats were placed individually in boxes for 5 min of acclimatization before testing, followed by a 5-minute rest period. The behavior of the rats was recorded for 15 min using a video camera (BASLER acA1280-60 g, USA) placed in the center above the box. LabVIEW software was used to capture the video images. MATLAB software was used to analyze the movement trajectories of the rats as described previously reference 54(Samson et al., 2015), and the total distance (cm), exercise time (s), and center zone rest time (s) were calculated.

### Modified version of the Y-maze test

The modified Y-maze test (Da et al., 2020; Da & Choi, 2020b) was used to assess working memory and behavioral flexibility. The apparatus consisted of black polyvinyl chloride boards with three arms, each 40 cm long, 10 cm wide, and 30 cm high. As shown in Figure supplement 9, the rats were acclimated for 2 days prior to testing. The trial consisted of two phases. In the sample phase trial, rats were placed individually in a maze with one of the three arms closed and were allowed to acclimate to the other two arms for 10 min. The test phase was conducted 24 h after the end of the sample phase. The previously closed arm was opened and defined as the new arm. The behavior of the animals in the maze was recorded for 10 min with a video camera (BASLER acA1280-60 g, USA) in a quiet room. MATLAB was used to analyze the total distance (cm), movement time (s), and new arm dwell time (s) of each group of rats in the Y-maze.

### Y-maze spontaneous alternation test

Spontaneous alternation in the Y-maze (Da et al., 2020; Da & Choi, 2020b) was used to assess spatial working memory during exploratory activities. During the acclimation phase, rats were allowed to freely explore the three arms of the Y-shaped maze for 30 minutes, and the ends of the three arms were marked as “A,” “B,” and “C,” respectively. The testing phase started from the center of the maze 10 min after the end of the adaptation phase, and the rats were allowed to move freely in the maze for 8 min. SAP was defined as consecutive visits to three different arms (ABC, ACB, BCA, and BAC). AAR indicated alternate arm return, and SAR indicated same arm return. The percentage of spontaneous alternation was calculated as follows: [(number of alternations)/ (total number of arms-2)]. It was used to assess the spontaneous exploratory behavior of the animals.

### T-maze, delayed non-match to place version

The T-maze test (Da et al., 2020; Da & Choi, 2020a) was used to evaluate the spatial memory ability of the animals. The experiment was carried out in a self-made T-maze made of polyvinyl chloride board; each arm was 40 cm long, 10 cm wide, and 30 cm high. In the adaptation stage, a small vessel was placed at the ends of the left and right arms of the T-maze, and 50 mg of food was placed inside. The experimental mice moved freely in the T-maze, and the experiment was started after 2 days of adaptation to foraging. The test was divided into two phases: sample and test. During the sampling stage, one arm of the maze was randomly selected, closed with a cardboard, and the experimental rat could only be forced to enter the open arm, take out the food after eating, and put it back into the cage. In the testing phase, the two arms were opened simultaneously; the arm with food in the sample phase was the wrong arm, and the other arm was the correct arm with the food reward. If the limbs of the rat enter any arm, the selection is considered valid, and no further backing is allowed. Each experimental mouse was tested seven times a day, with an interval of 15 min between each test. The spatial memory ability of animals was evaluated based on the correct rate of rat selection.

### Spike sorting and firing rate analysis

With 80 μV as the threshold, high-pass signals (> 300 Hz) with trough values above the threshold were considered spikes. Spikes were extracted by principal component analysis (Jing, Si, Olson, & He, 2005) and K-means clustering methods (Chah et al., 2011) in BlackRock Offline Spike Sorter (BOSS) software (1.2.0, Blackrock, USA), and were manually combined, divided, or dropped to refine individual neuron clusters based on autocorrelation mid-refractory periods (ISI < 2 ms). The average firing frequency per second within a single neuron spike was calculated.

### Relative power and sample entropy

LFP data were imported into MATLAB (MathWorks Inc., Natick, MA) using built-in and custom programs to calculate the absolute power and sample entropy of the theta (4–8 Hz) band, beta (15–30 Hz) band, and gamma (30–45 Hz) band. The EEG signals of all the above frequency bands were Fourier transformed to obtain the absolute power of each frequency band, and the total absolute power of the frequency band was obtained by adding the values of each band. The relative power of each band is equal to the corresponding absolute power divided by the total absolute power. The sample entropy of the above frequency bands was calculated simultaneously based on a previous study (Richman & Moorman, 2000). The sample entropy was calculated using the following equation:

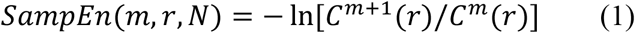

where N is the length of the data, m is the vector dimension, and r is tolerance. The length of each data segment was selected to be 1000; vector dimension, 2; and tolerance, 0.2. After each data segment was calculated, the sample entropy of each group was calculated using an averaging method.

### Western blotting

The rats were anesthetized by intraperitoneal injection of a chloral hydrate solution. After decapitation, brain tissue rapidly peeled off. The prefrontal lobe tissues were then harvested. The tissue samples were placed in a tube, and a RIPA lysis solution was added. After mixing, tissues were lysed at 4°C overnight and then centrifuged at 12,000 *g* for 10 min at 4°C. The supernatant was then collected and stored for later use. The sample was denatured at 100°C for 10 min, equilibrated at room temperature, centrifuged at 10,000*g* for 5 min, and set aside. The sample solution was transferred to a membrane after sodium dodecyl-sulfate polyacrylamide gel electrophoresis and blocked with a blocking solution at 37°C for 2 h. After blocking, the membrane was washed and incubated with primary antibodies (mBDNF (NB100-98682, Novus Biologicals, LLC., Centennial, CO, USA) and DAT (AB1591P, Millipore Corporation, Billerica, MA, USA)) overnight at 4°C. After rinsing three times with phosphate-buffered saline/Tween (10 min each), the secondary antibody (Zhongshan Jinqiao, Beijing, ZB2301) was incubated at room temperature for 90 min. Quantification of band immunoreactivity was performed using an enhanced chemiluminescence system (Pierce, Rockford, IL, USA) and was imaged using an Image Quant LAS-4000 imaging system (Fujifilm, Tokyo, Japan). Finally, the target band on the gel was scanned using a computer, and grayscale analysis was performed using Quantity one software. The protein expression/glyceraldehyde 3-phosphate dehydrogenase ratio was then calculated.

## Statistical Analysis

Data are presented as the mean ± standard error of the mean (SEM). Data were analyzed by the Kruskal–Wallis test. All statistical analyses were performed using MATLAB (MathWorks Inc., MA, USA), and p < 0.05 was considered statistically significant.

## CRediT authorship contribution statement

Mengran Wang: Methodology, Software, Data curation, Writing –original draft. Teng Wang: Methodology, Software, Data curation. Hui Ji: Methodology. Jiaqing Yan: Methodology, Software. Xingran Wang: Data curation, Writing–original draft. Xiangjian Zhang: Writing– original draft. Xin Li: Writing –review & editing. Yi Yuan: Supervision, Conceptualization, Methodology, Software, Writing –review & editing.

## Declaration of Competing Interest

The author(s) declare no potential conflicts of interest with respect to the research, authorship, and/or publication of this article.

## Data and code availability statement

The data and code that support the findings of this study are available from the corresponding authors, upon reasonable request.

## Ethics statement

All procedures were conducted according to the guidelines of the Animal Ethics and Administrative Council of Yanshan University.

## Acknowledgments

This research was supported by National Natural Science Foundation of China (62273291), Natural Science Foundation of Hebei Province (F2022203050), National Natural Science Foundation of China (61827811), Scientific and Technological Innovation 2030 (No. 2021ZD0204300).

## Supplementary materials

**Figure supplement 1.**
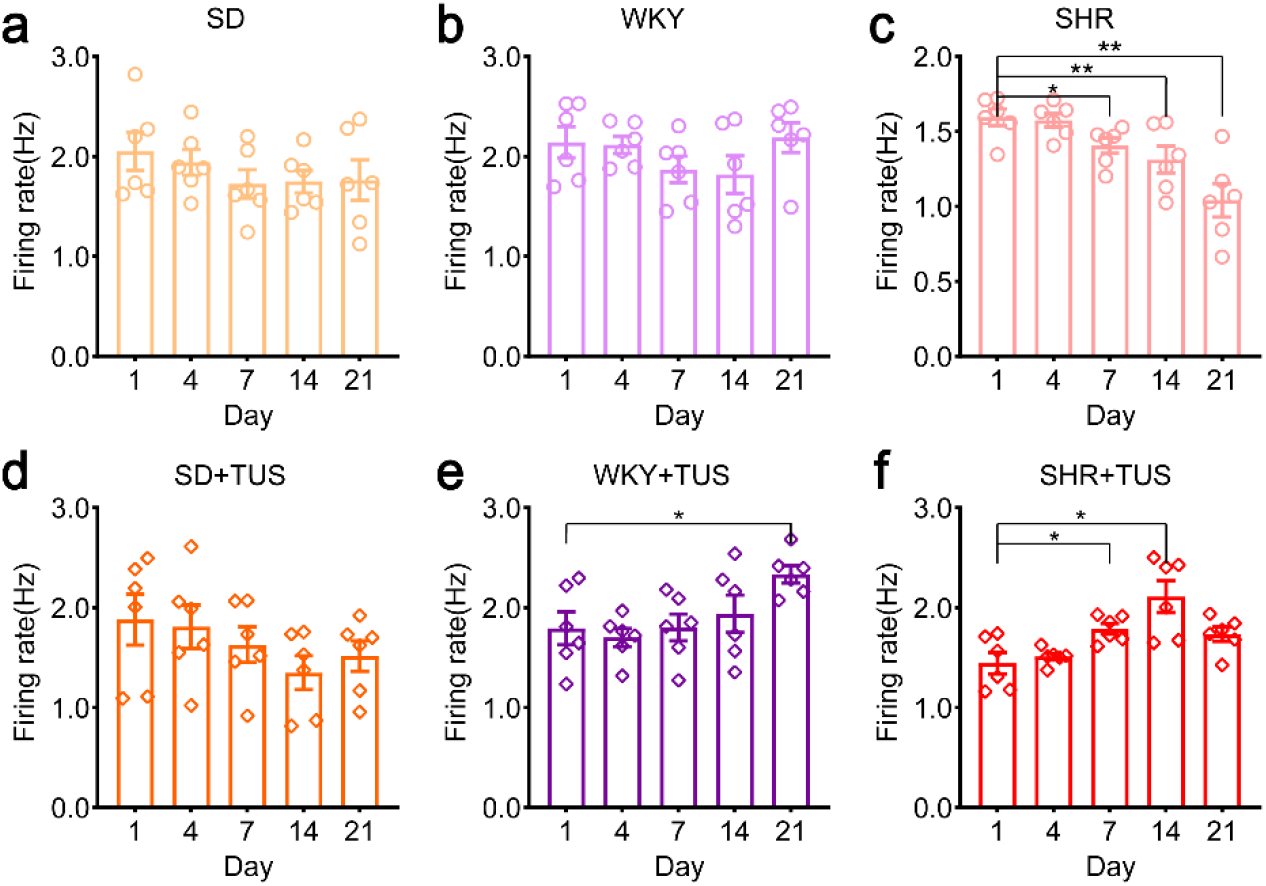
Average of spike firing rate in different groups. (a) SD group. (b) SD+TUS group. (c) WKY group. (d) WKY+TUS group. (e) SHR group. (f) SHR+TUS group. (N=6 for each group, Mean±SEM, *p <0.05, **p<0.01, Kruskal–Wallis test).

**Figure supplement 2.**
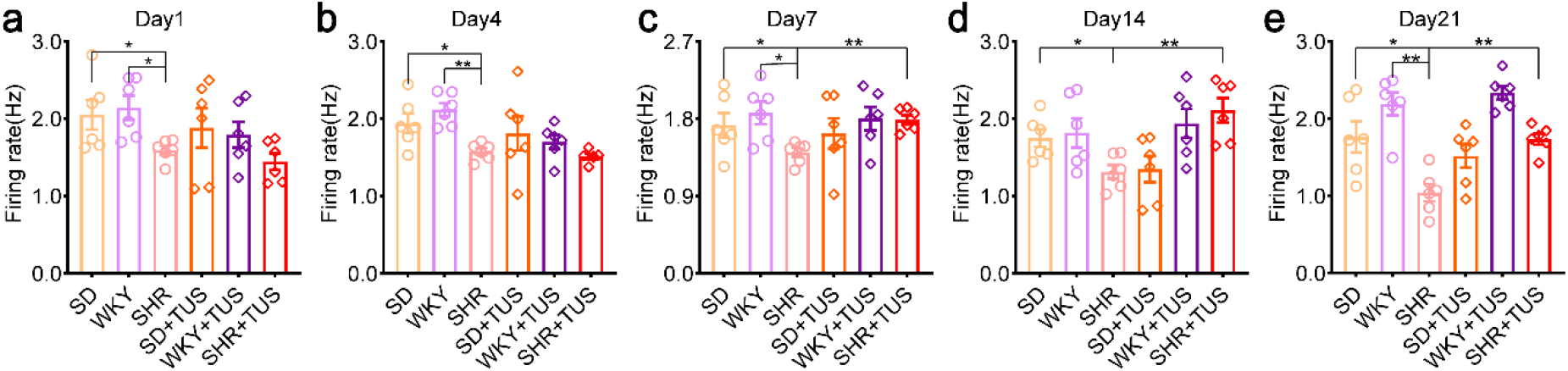
Average of spike firing rate in 1,4,7,14 and 21 day after TUS. (a) day 1. (b) day 4. (c) day 7. (d) day 14. (e) day 21. (N=6 for each group, Mean±SEM, *p <0.05, **p<0.01, Kruskal–Wallis test).

**Figure supplement 3.**
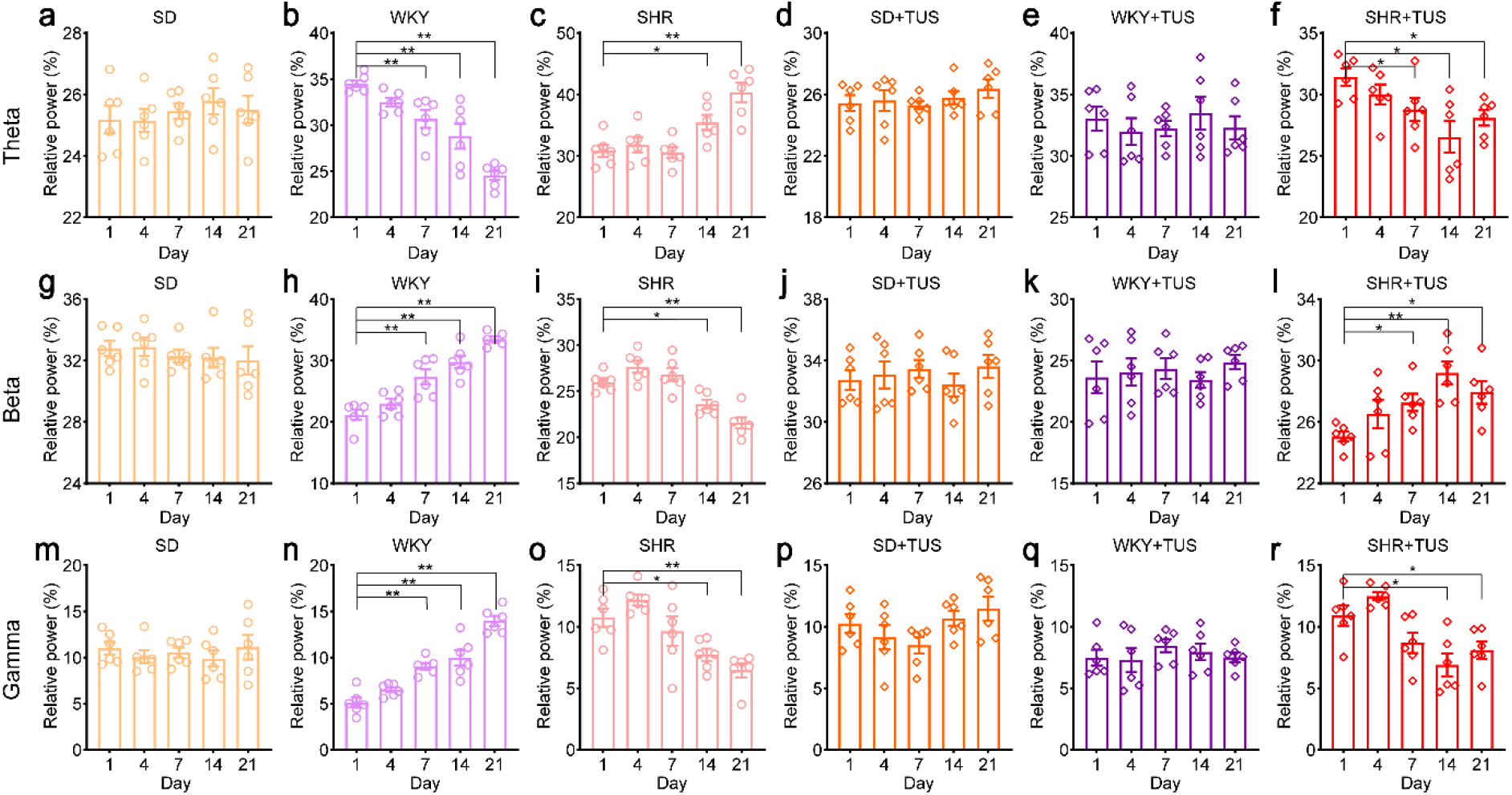
The relative power analysis of LFPs at the different frequency bands in different groups. (a)-(f) The relative power analysis of LFPs at the theta (4–8 Hz) frequency band in different groups, (a) SD group, (b) WKY group, (c) SHR group, (d) SD+TUS group, (e) WKY+TUS group, (f) SHR+TUS group. (g)-(l) The relative power analysis of LFPs at the beta (13-30 Hz) frequency band in different groups (g) SD group, (h) WKY group, (i) SHR group, (j) SD+TUS group, (k) WKY+TUS group, (l) SHR+TUS group. (m)-(r) The relative power analysis of LFPs at the gamma (30-45 Hz) frequency band in different groups, (m) SD group, (n) WKY group, (o) SHR group, (p) SD+TUS group, (q) WKY+TUS group. (r) SHR+TUS group. (N=6 for each group, Mean±SEM, *p <0.05, **p<0.01, Kruskal–Wallis test).

**Figure supplement 4.**
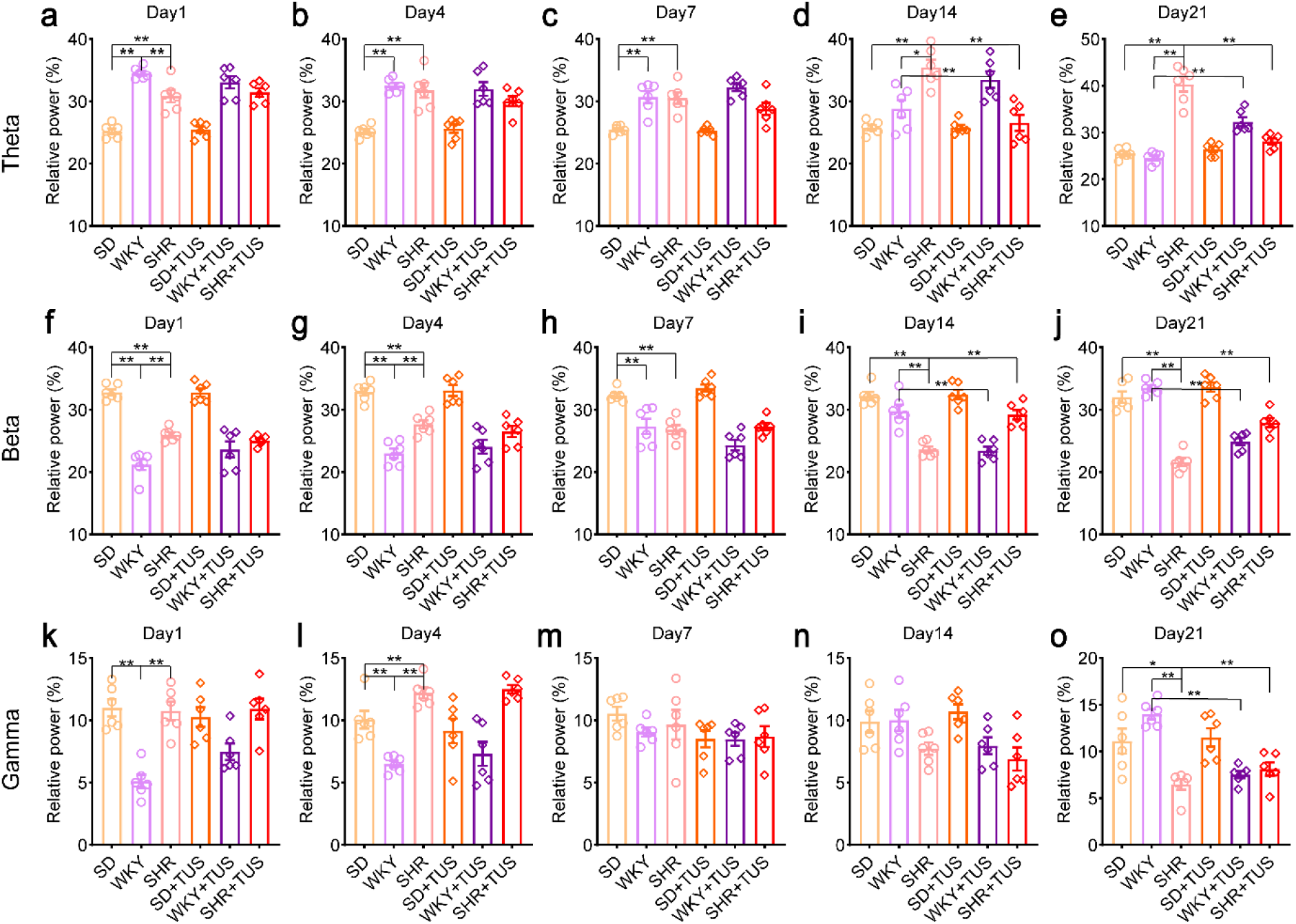
The relative power of LFPs at the different frequency bands on 1,4,7,14 and 21 day after TUS. The relative power of LFPs at the theta (4-8Hz) frequency band in different groups at (a) day 1, (b) day 4, (c) day 7, (d) day 14, (e) day 21. The relative power analysis of LFPs at the beta (13-30Hz) frequency band in different groups at (f) day 1, (g) day 4, (h) day 7, (i) day 14, (j) day 21. (k) The relative power of LFPs at the gamma (30-45Hz) frequency band in different groups at day 1, (l) day 4, (m) day 7, (n) day 14, (o) day 21. (N=6 for each group, Mean±SEM, *p <0.05, **p<0.01, Kruskal–Wallis test).

**Figure supplement 5.**
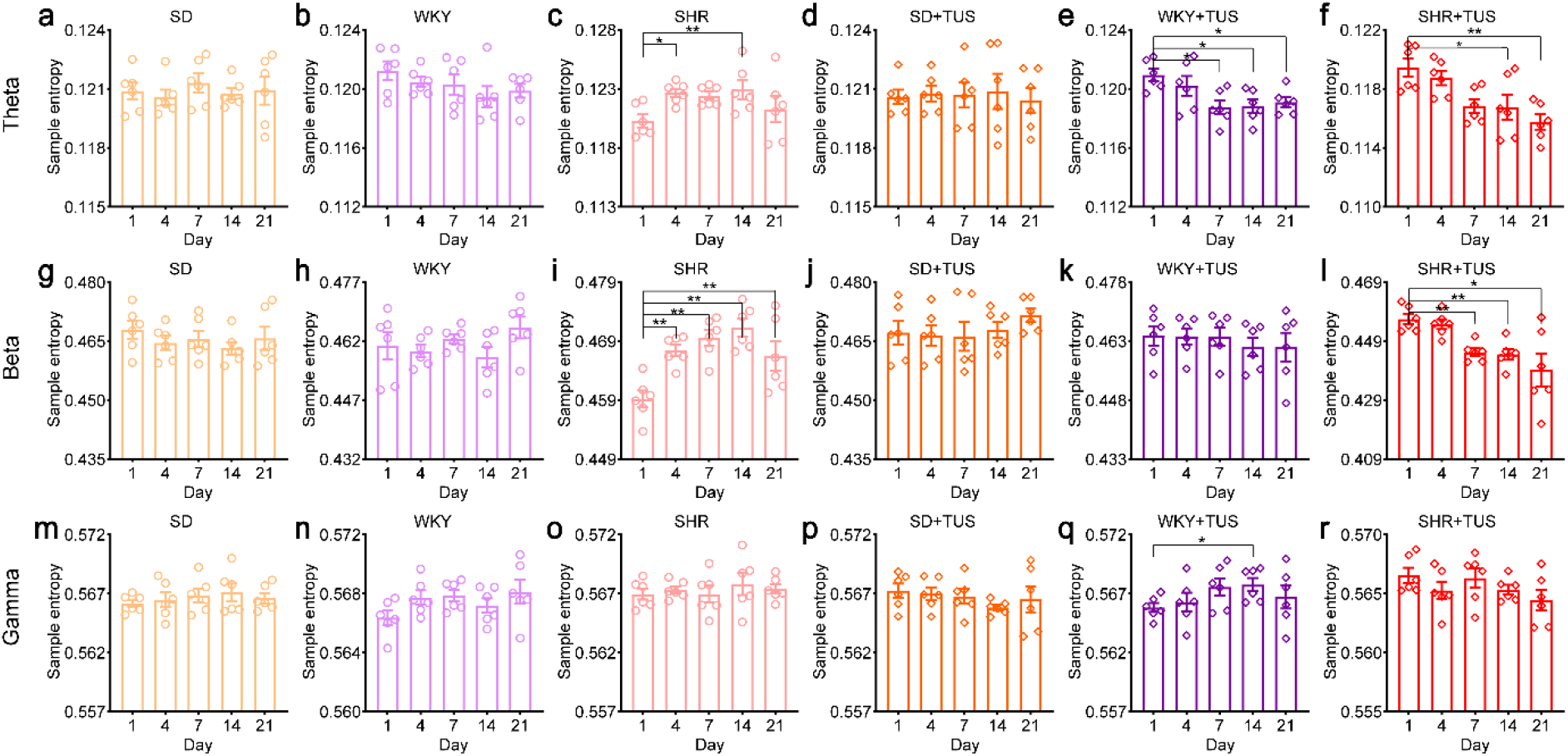
The sample entropy of LFPs at the different frequency bands in different groups. (a)-(f) The sample entropy of LFPs at the theta (4–8 Hz) frequency band in different groups, (a) SD group, (b) WKY group, (c) SHR group, (d) SD+TUS group, (e) WKY+TUS group, (f) SHR+TUS group. (g)-(l) The sample entropy of LFPs at the beta (13-30 Hz) frequency band in different groups (g) SD group, (h) WKY group, (i) SHR group, (j) SD+TUS group, (k) WKY+TUS group, (l) SHR+TUS group. (m)-(r) The sample entropy of LFPs at the gamma (30-45 Hz) frequency band in different groups, (m) SD group, (n) WKY group, (o) SHR group, (p) SD+TUS group, (q) WKY+TUS group. (r) SHR+TUS group. (N=6 for each group, Mean±SEM, *p <0.05, **p<0.01, Kruskal–Wallis test).

**Figure supplement 6.**
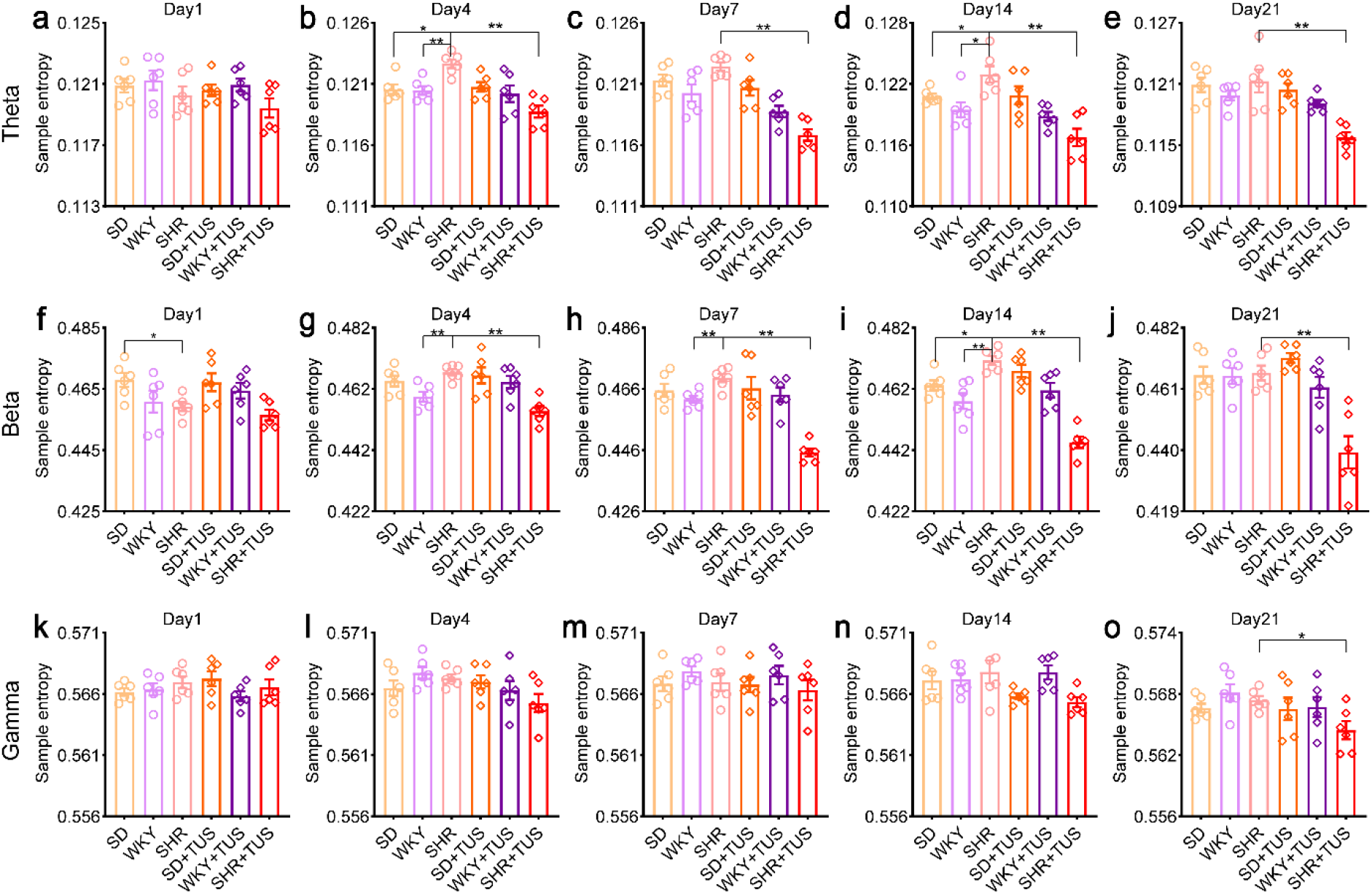
The sample entropy of LFPs at the different frequency bands on 1,4,7,14 and 21 day after TUS. The sample entropy of LFPs at the theta (4-8Hz) frequency band in different groups at (a) day 1, (b) day 4, (c) day 7, (d) day 14, (e) day 21. The sample entropy of LFPs at the beta (13-30Hz) frequency band in different groups at (f) day 1, (g) day 4, (h) day 7,(i) day 14, (j) day 21. (k) The sample entropy of LFPs at the gamma (30-45Hz) frequency band in different groups at day 1, (l) day 4, (m) day 7, (n) day 14, (o) day 21. (N=6 for each group, Mean±SEM, *p <0.05, **p<0.01, Kruskal–Wallis test).

**Figure supplement 7.**
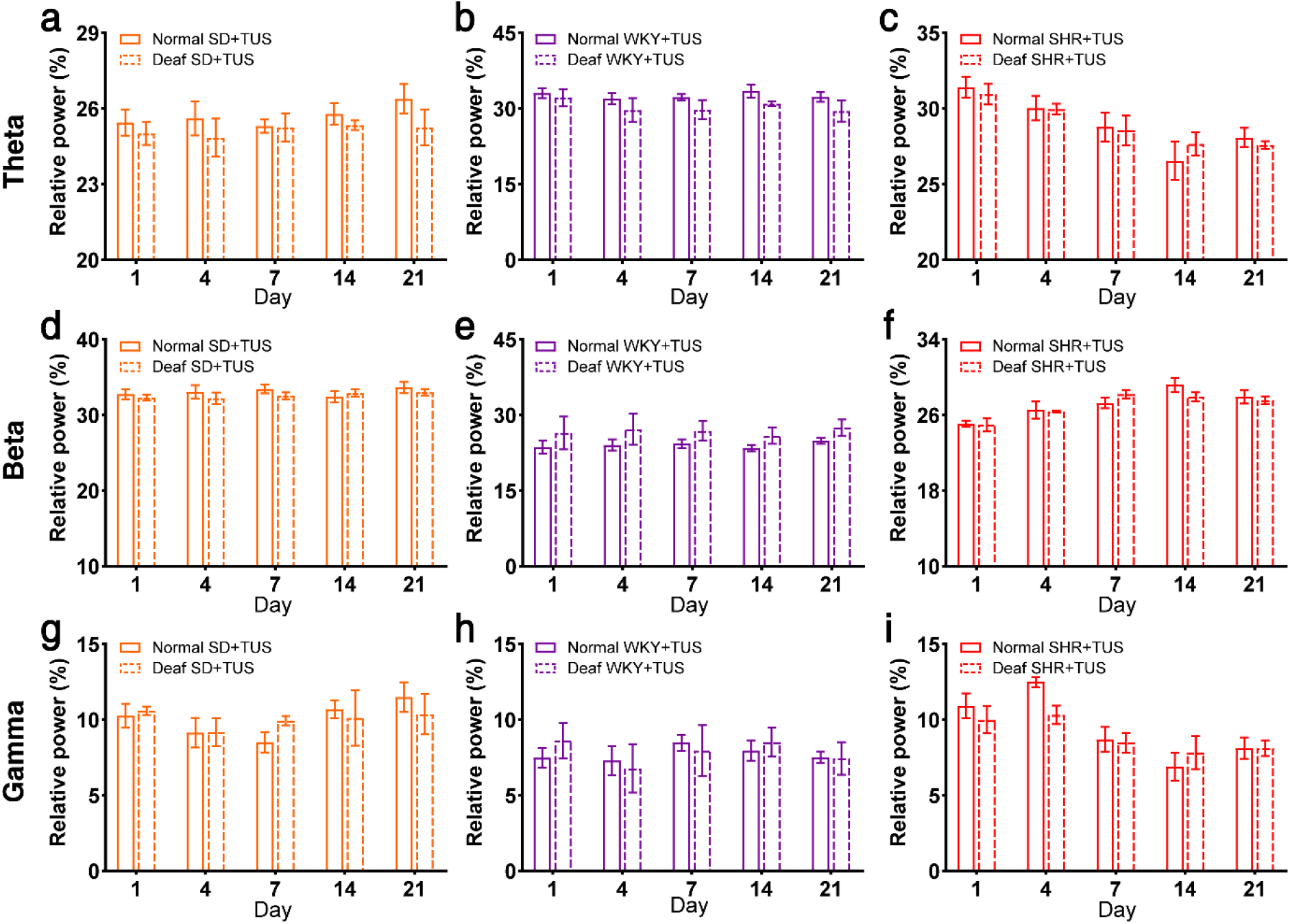
Comparisons of the relative power of LFPs between normal and deaf SD+TUS, WKY+TUS, SHR+TUS groups at the different frequency bands. (a)-(c)The relative power of LFPs at the theta frequency bands in normal and deaf SD+TUS, WKY+TUS, SHR+TUS groups, (a) normal and deaf SD+TUS groups, (b) normal and deaf WKY+TUS groups, (c) normal and deaf SHR+TUS groups. (d)-(f) The relative power of LFPs at the beta (13–30Hz) frequency bands in normal and deaf SD+TUS, WKY+TUS, SHR+TUS groups, (d) normal and deaf SD+TUS groups, (e) normal and deaf WKY+TUS groups, (f) normal and deaf SHR+TUS groups. (g)-(i) The relative power of LFPs at the gamma (30–45Hz) frequency bands in normal and deaf SD+TUS, WKY+TUS, SHR+TUS groups, (g) normal and deaf SD+TUS groups, (h) normal and deaf WKY+TUS groups, (i) normal and deaf SHR+TUS groups. (Mean±SEM, n=3 for each group, p>0.05, Kruskal–Wallis test).

**Figure supplement 8.**
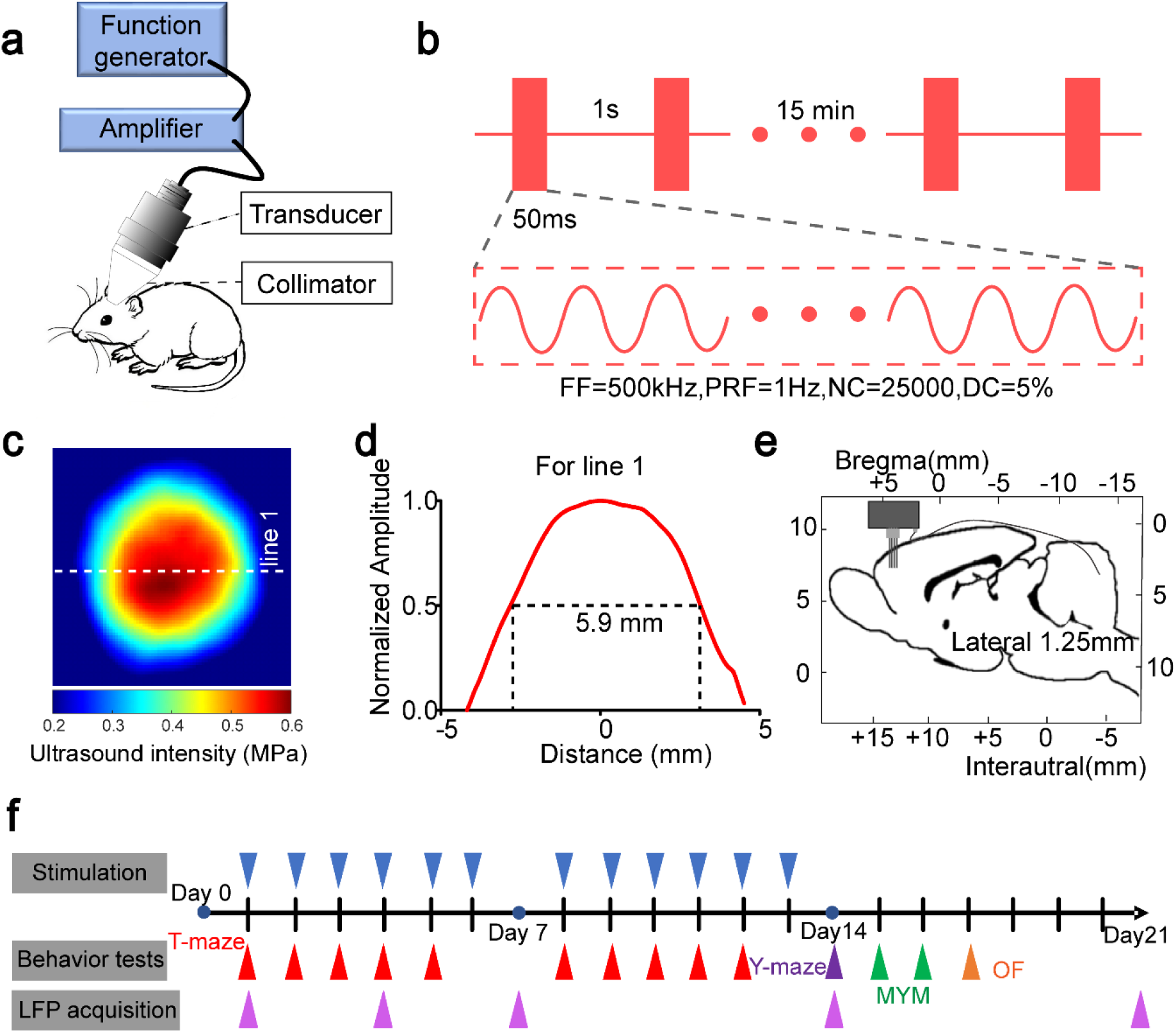
Experimental schematic diagram. (a) Schematic diagram of the TUS set-up. (b) Definition of the ultrasound parameters: duty cycle (DC)and fundamental frequency (FF). (c) Pseudocolor plots showing the ultrasound pressure field. (d) The reconstruction profile along the white dotted lines in Figure1(c). (e) Schematic diagram showing the electrode position and corresponding cerebral cortical region. (f) The time schedule of TUS, behavior tests and Spikes and LFP acquisition. OF, open-field test; MYM, modified version of the Y-maze test.

**Figure supplement 9.**
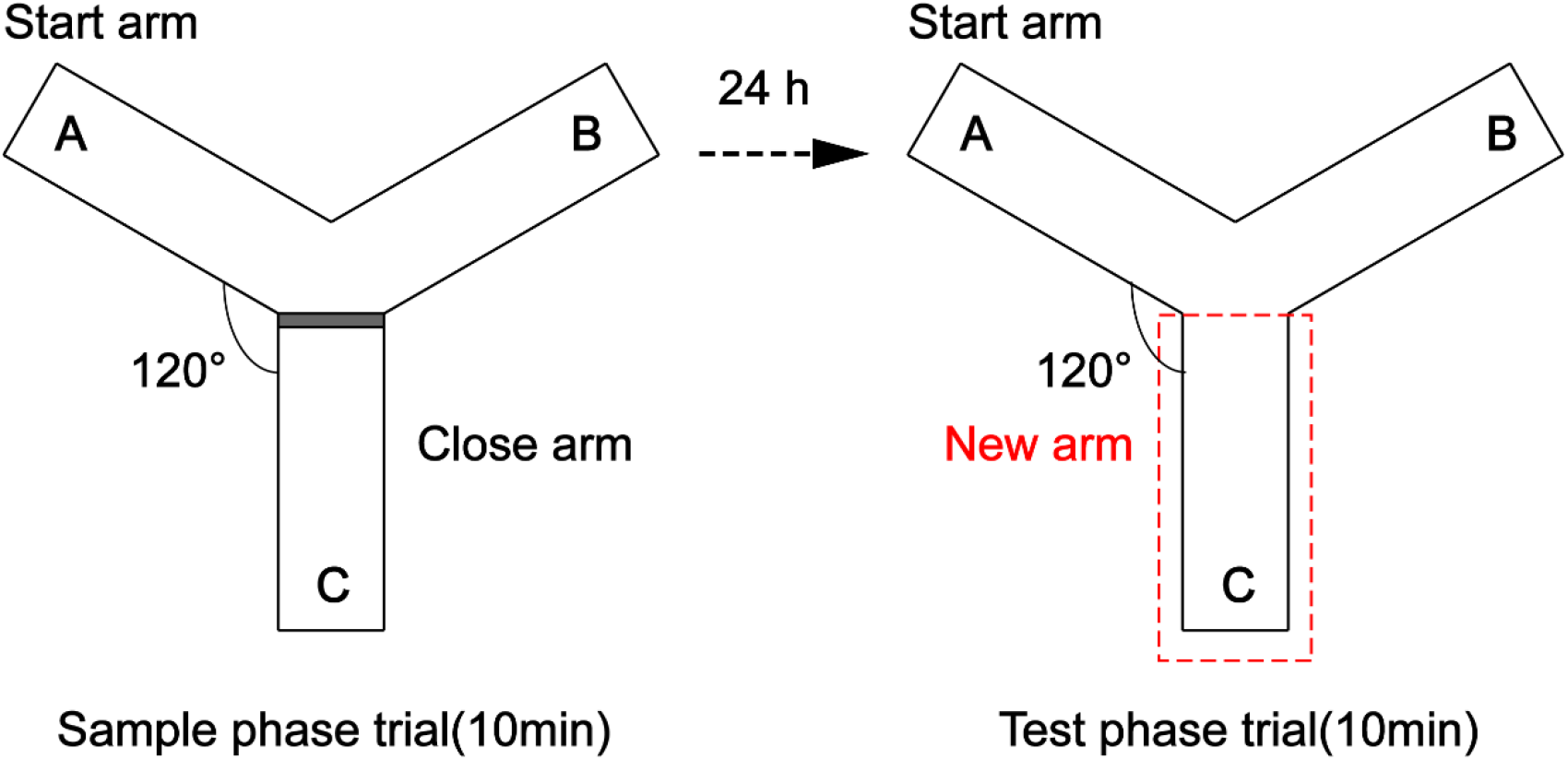
Schematic of modified Y-maze.

## Notes

### Competing Interest Statement

The authors have declared no competing interest.

